# BAGEL: A computational framework for identifying essential genes from pooled library screens

**DOI:** 10.1101/033068

**Authors:** Traver Hart, Jason Moffat

## Abstract

**Background:** The adaptation of the CRISPR-Cas9 system to pooled library gene knockout screens in mammalian cells represents a major technological leap over RNA interference, the prior state of the art. New methods for analyzing the data and evaluating results are needed.

**Results:** We offer BAGEL (Bayesian Analysis of Gene EssentiaLity), a supervised learning method for analyzing gene knockout screens. Coupled with gold-standard reference sets of essential and nonessential genes, BAGEL offers significantly greater sensitivity than current methods, while computational optimizations reduce runtime by an order of magnitude.

**Conclusions:** Using BAGEL, we identify ~2,000 fitness genes in pooled library knockout screens in human cell lines at 5% FDR, a major advance over competing platforms. BAGEL shows high sensitivity and specificity even across screens with highly variable reagent quality.

## Background

Perturbing gene activity and evaluating the resulting phenotype is a fundamental technique for identifying the biological processes in which a gene participates. Traditionally, the ability to induce complete gene knockouts on a genomic scale has been exclusively the domain of model organisms such as yeast, while experiments in higher eukaryotes, including human cell lines, have relied on RNA interference (RNAi). RNAi uses the endogenous RNA-induced silencing complex (RISC) machinery to target messenger RNA transcripts, which have a very large dynamic range of abundance, resulting in data that is often diluted by incomplete target knockdown and off-target effects of variable severity [1–3].

The adaptation of CRISPR-Cas9 technology to pooled library gene knockout screens in mammalian cells allows the identification of genes whose knockout contributes to gene fitness [4–8]. A pooled library screen typically contains several guide RNAs (gRNA) targeting each gene, and large numbers of cells are treated such that each cell is affected by (on average) a single gRNA clone, while each gRNA species targets hundreds of cells. Unperturbed cells, or cells with knockouts showing no growth phenotype, grow at wildtype rates, while cells harboring a guide RNA that targets a gene resulting in a growth defect show lower growth rates (Figure 1A). To identify the genes whose knockout causes a fitness defect, the frequency distribution of gRNA in the population is assayed by deep sequencing and compared to the frequency distribution at an early control timepoint. Changes in the frequency distribution of gRNA are measured as log fold changes (Figure 1A, sidebar) where severe negative fold changes reflect gRNA that cause severe fitness defects.

**Figure 1.**
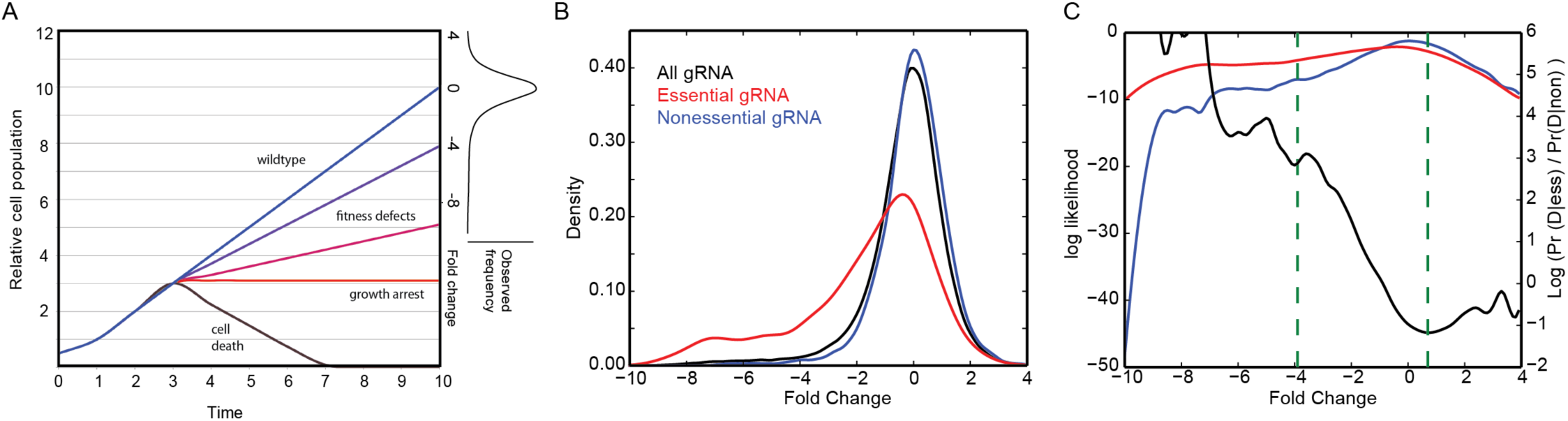
BAGEL overview. (A) Simulated growth curves of wildtype cells (blue), which double at every time increment. When genetic perturbations are induced (T=3), moderate (purple) to severe (magenta) fitness defects, growth arrest (red), and cell death (black) result in different relative growth rates. At sampled timepoints, fold change relative to wildtype growth is the readout from a sequencing assay. (B) Representative data from one replicate. The fold change distribution of all gRNA targeting essential genes (red) is shifted relative to the fold change distribution of all gRNA targeting nonessential genes (blue). The fold change distribution for all gRNA (black) is shown for reference. (C) The log likelihood functions of the red and blue curves from (B), left Y axis. The BAGEL method calculates the log likelihood ratio (black, right Y axis) of these two curves, within empirical boundaries (green dashes), for each bootstrap iteration; see Methods for details.

Aggregating individual reagent effects into an accurate estimate of gene-level effect is a major challenge in the analysis of pooled library screen data[9–13]. To analyze pooled library RNAi screens, which have similar experimental design, we developed an early Bayesian classifier and demonstrated its superiority over contemporary approaches [2]. A key feature of this study was the establishment of reference sets of core essential and nonessential genes, which can be used as gold standards to evaluate other algorithms in analyzing fitness screens. Here we describe BAGEL, the Bayesian Analysis of Gene EssentiaLity, an adaptation of the previously described Bayesian classifier. BAGEL features a more robust statistical model, major performance enhancements, and an improved user interface. BAGEL source code, documentation, and reference files are available at http://bagel-for-knockout-screens.sourceforge.net/.

## Methods

A pooled library CRISPR-Cas9 fitness screen in human cells involves having multiple gRNA reagents targeting each gene and is often evaluated at several timepoints, ideally with multiple replicates at each timepoint. BAGEL first estimates the distribution of fold changes of all gRNA targeting all genes in either the essential or nonessential training sets (Figure 1B). Then, for each withheld gene, it evaluates the likelihood that the observed fold changes for gRNA targeting the gene were drawn from either the essential or the nonessential training distributions. The result is a Bayes Factor:

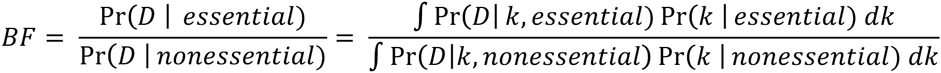
where the data, D, is the set of observed fold changes for the uncharacterized gene and *k* is the fold change distribution of the training set, empirically estimated using a kernel density estimate function (Figure 1B, red and blue curves).

The integral is estimated by bootstrap resampling of the training sets. At each iteration the *k* distributions are calculated and a log BF is calculated:

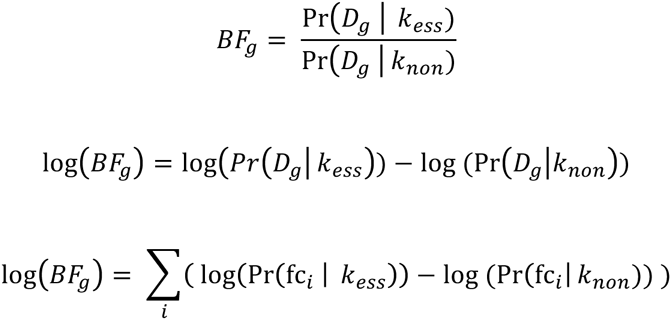
where fc*_i_*,· are the observed fold changes for gRNA targeting gene *g*. One thousand bootstrapping iterations are conducted; a Bayes Factor is calculated for each iteration and the mean and standard deviation of the resulting posterior distribution of BFs is reported.

Two factors inherent in the data require that empirical boundaries be applied to the calculations. First, when taking the ratio of two curves, the ratio can take on extreme values when the denominator approaches zero. Second, kernel density estimates become unstable in regions of sparse data. For these reasons, we identify the lowest fold change (x-coordinate) at which the k_non_ density estimate, the denominator above, exceeds 2^−7^, and set this as a lower bound (Figure 1C). All observed changes below this boundary are set to the boundary value. Similarly, we calculate the fold change at which the log ratio of the curves is a minimum and set this as an upper bound (Figure 1C). These boundaries ensure that individual observations do not dominate the final BF score while, in our experience, making no material change to gene estimates: observed fold changes outside these boundaries are not stronger evidence that a gene does or doesn’t induce a fitness defect, given the normal constraints of the experiment (number of cells, sequencing depth, etc.). Note that this approach makes no statement about whether a gene knockout can increase cell fitness, only whether perturbation causes a growth defect.

For very large CRISPR libraries, the calculation as described can be computationally expensive. To speed up the calculations, we include two optimizations. First, we round all calculated fold changes to the nearest 0.01. Second, for each bootstrap iteration, we calculate the value of the log ratio function (Figure 1C) at each 0.01 within the empirical boundaries described above and store the values in a lookup table. Then, instead of recalculating the values for each gRNA, we pull the value of the log ratio function from the lookup table. These optimizations decrease processing time by over an order of magnitude, with no impact on final results (Pearson’s r ~ 0.999 for final BFs; data not shown).

For knockout screens with multiple timepoints, the BF is calculated at each timepoint, and a final BF is the sum of the timepoint BFs. Since the posterior BF distributions are approximately normal (by KS tests, not shown), the variance of the final BF is estimated as the sum of the variances at the timepoints.

## Results and discussion

We demonstrate this approach with screens from the Toronto KnockOut (TKO) library in four cell lines: a patient-derived glioblastoma cell line (GBM, Figure 2A), HCT116 colorectal carcinoma cell line (Figure 2B), HeLa cervical carcinoma cell line (Figure 2C), and RPE1 retinal pigmented epithelial cells (Figure 2D) [14]. All the screens were sampled at multiple timepoints. Using the gold-standard reference sets from [2], BFs were calculated for each timepoint and precision-recall (PR) curves were plotted. In all cases, later timepoints showed improved recall over the earliest timepoint. The “integrated” sample is the sum of the timepoint BFs and can be considered a summary result for the entire screen; the PR curve for the integrated sample is in every case as good or better than the timepoint curves. In all cases screens yielded a very large number of fitness genes: on average, ~2,000 genes at 5% false discovery rate (FDR) using the integrated results, and these genes show very high functional coherence (see [14] for a more complete evaluation).

**Figure 2.**
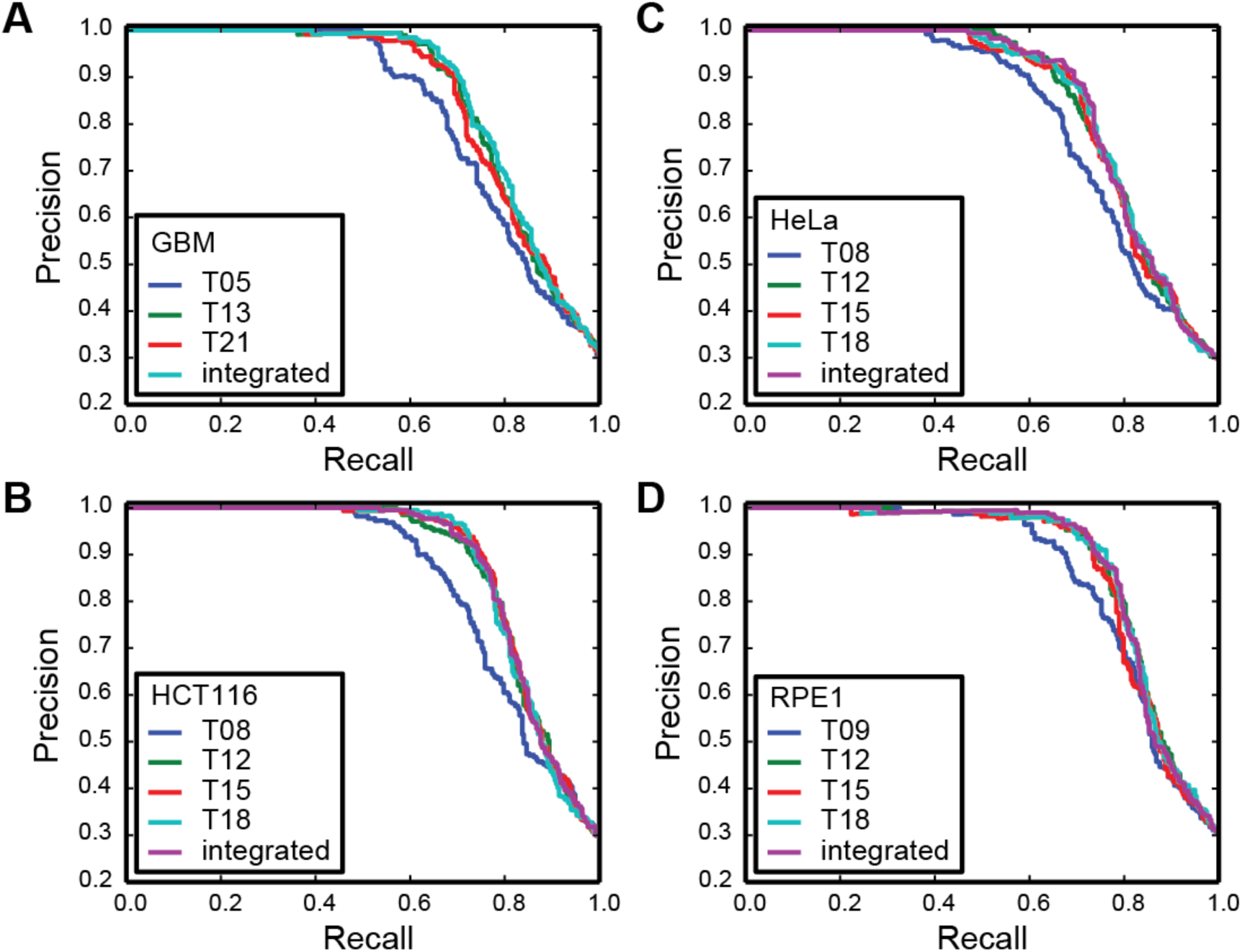
**Precision-recall curves for BAGEL results** for GBM (A), HCT116(B), HeLa (C), and RPE1 (D) screens using the TKO library. Where indicated, a single timepoint is plotted. “Integrated” = Bayes Factors summed across all timepoints in the experiment.

One question that arises from these results is whether the lower performance at the early timepoint, relative to the later ones, reflects the screening technology or the biology of the systems being perturbed. We address this question by looking at functional enrichment in genes unique to the early hits, genes unique to the late hits, or genes in the intersection, using the GORILLA web service[15]. We find that most (75–89%) early hits are also observed at the last timepoint (Figure 3A), and that genes exclusively in the early hit set are not meaningfully enriched for annotated biological processes. Looking specifically at the GBM cell line, genes in the intersection are highly enriched for core biological processes one could reasonably expect to cause fitness defects (Figure 3B). Genes in the intersection comprise only 53–65% of the total number of hits at the last timepoint; however, genes exclusive to the last timepoint typically extend coverage of the biological processes identified in the intersection, as shown for GBM cells in Figure 3B, and identify few novel processes.

**Figure 3.**
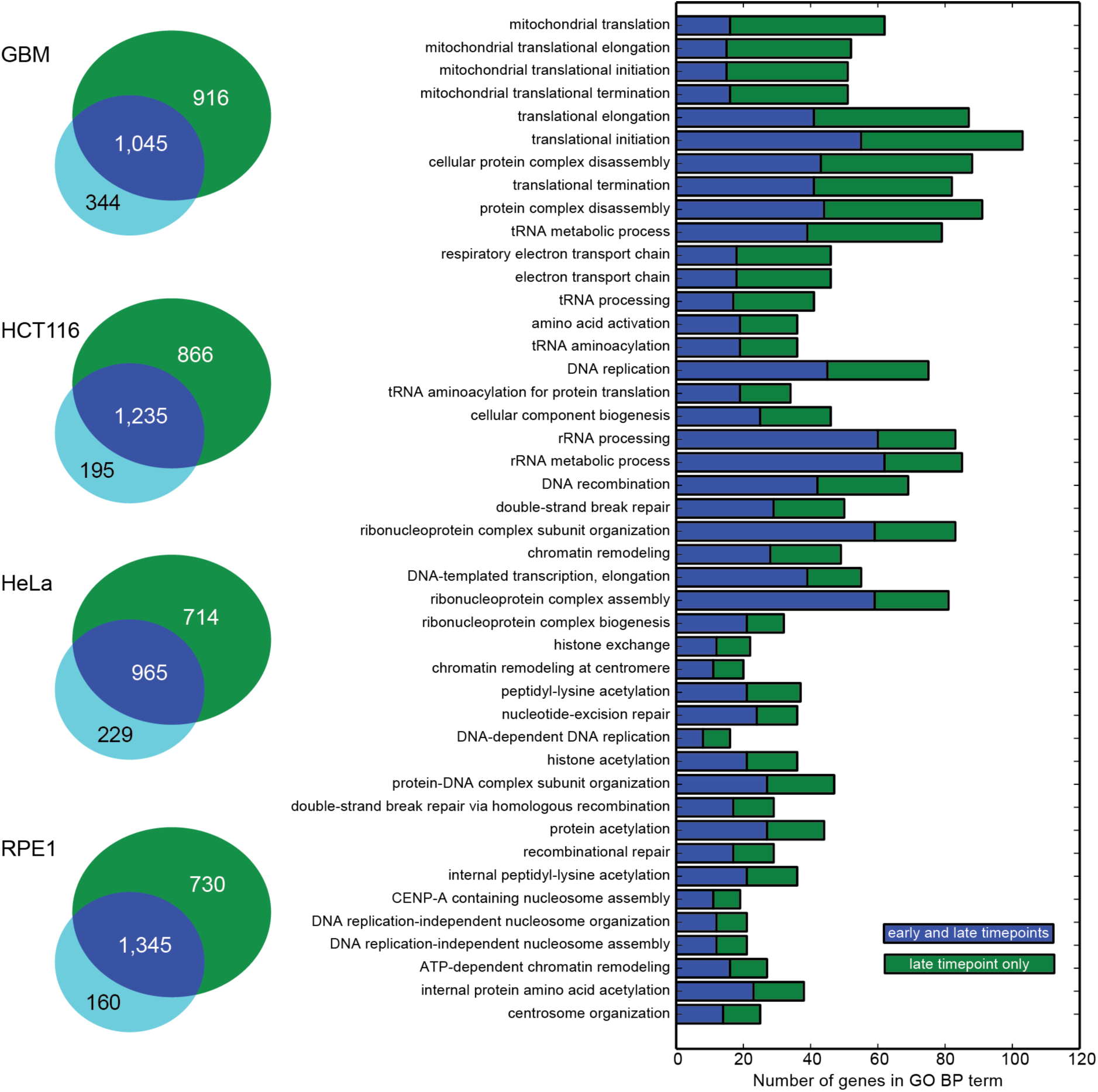
Comparing early and late hits. (A) Number of fitness genes detected at early timepoint (cyan), late timepoint, (green), or both (blue) in each TKO screen. (B) Representative data from GBM screen. Most GO_BP terms enriched in late-only genes (green) extend observations of terms enriched in genes found in both early and late timepoints.

Though late fitness genes typically reflect the processes observed in early fitness genes, genes which encode proteins involved in mitochondrial function offer an interesting contrast. Genes in both the early and late timepoints are enriched for some mitochondrial processes, including protein transport to the mitochondrion and mitochondrial translation. However, the late-only genes are enriched for a small number of GO BP terms that are centered around functions related to oxidative phosphorylation, including “respiratory chain complex I assembly” (7 hits of 18 annotated genes, 7.4-fold enrichment), “respiratory chain complex IV assembly” (4/8 genes, 9.4-fold), and “mitochondrial electron transport, NADH to ubiquinone” (12/36 genes, 6.3-fold). This difference may reflect a more subtle phenotype (i.e. lower fitness defect) among oxphos genes that only becomes detectable at the later timepoint (Figure 1A).

We compared BAGEL to MAGeCK, a contemporary method for analyzing CRISPR knockout screens[10]. MAGeCK ranks gRNAs by P-value derived from a negative binomial model comparing control to experimental timepoints, then calculates gene-level P-values using modified Robust Ranking Aggregation. We compared MAGeCK results to BAGEL results using only the final timepoint from the TKO screens described above, and plotted PR curves using the same reference sets (Figure 4A-D). In all cases, BAGEL outperformed MAGECK, yielding more recall and more overall hits in a reasonable range of empirically-calculated FDR (5–15%). Most striking, however, was the severe lack of sensitivity using the theoretical model of MAGeCK. Although gene rankings for the two methods were generally similar (Spearman correlations 0.76–0.81 for the top 3,000 genes in each set), the MAGeCK algorithm yielded only 674 (mean; range 489–905) genes at 10% FDR, using its own FDR estimates (Figure 4). We are confident that the higher numbers of fitness genes detected by BAGEL are in fact real: we analyze their expression level, biological function, and other functional genomic data in detail in [14].

**Figure 4.**
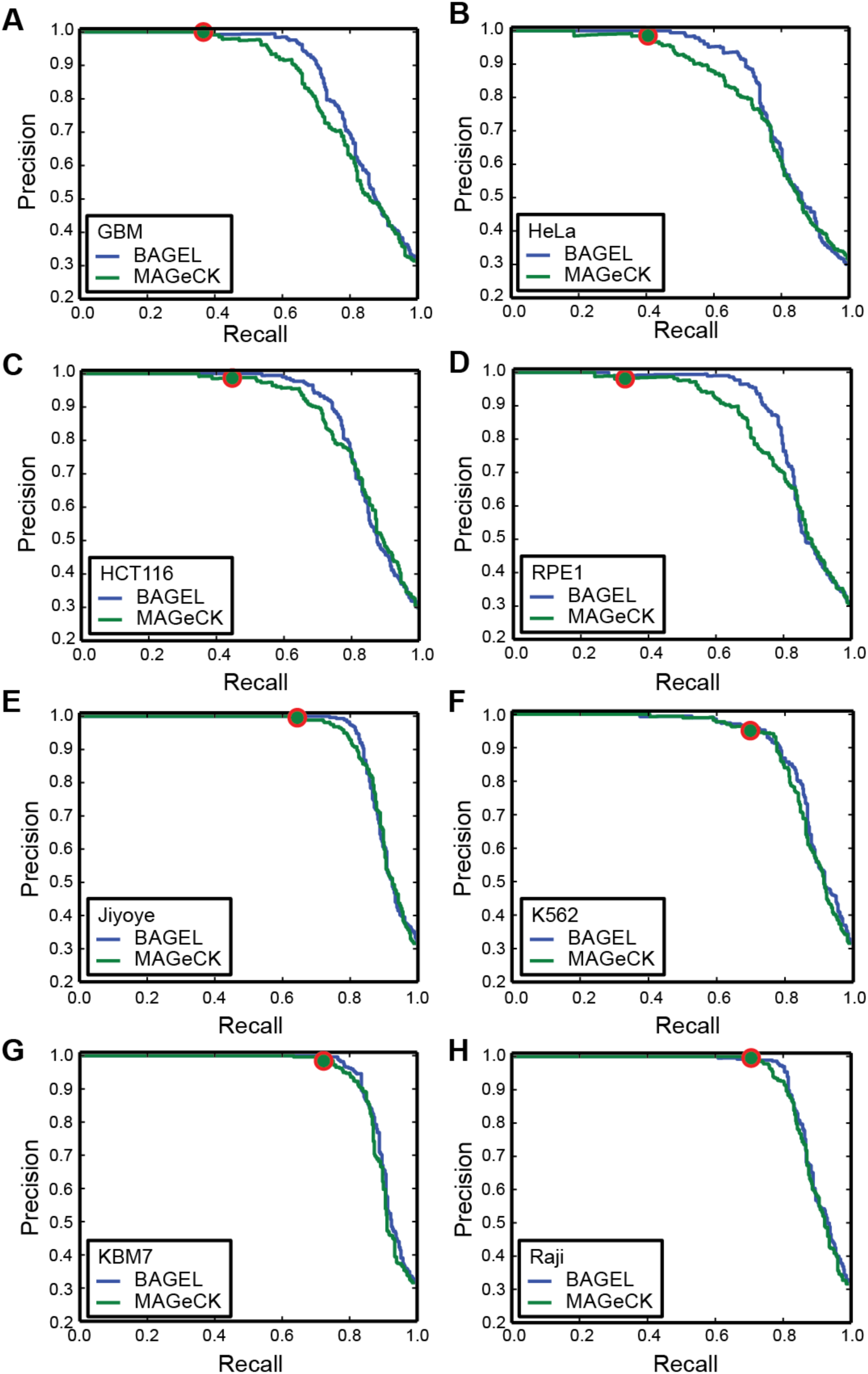
Comparing BAGEL with MAGeCK. For each cell line, precision-recall curves were plotted for BAGEL and MAGeCK results using the last timepoint of the screen. Red circle indicates results at MAGeCK-reported 10% FDR cutoff. (A-D) TKO screens from Hart *et al*. [14] (E-H) Screens from Wang *et al*. [16]

We also compared the two algorithms using a newly published data set from Wang *et al*. [16], where four leukemia and lymphoma cell lines were screened for essential genes using a large gRNA library. As with the TKO screens, the BAGEL algorithm yields equal or superior precision-recall curves and greater sensitivity, though with a smaller margin of improvement (Figure 4E-H). MAGeCK identifies 1,571 (mean, range 1,241–1,800) hits at 10% FDR while BAGEL identifies on average 2,272 (range 1,963–2,482) essential genes at 5% FDR.

The reason behind the difference in sensitivity between BAGEL and MAGeCK likely lies in the variable effectiveness of CRISPR reagents. Examining the fold change distribution of all guides targeting genes in the reference set of high-confidence essentials (Figure 1B), it is evident that many gRNAs targeting essential genes do not show significant dropout. The BAGEL algorithm chooses between the essential and nonessential distributions, and is able to detect even a slight shift in overall effect, whereas a statistical test based solely on excluding the null hypothesis – generally speaking, that the observed fold changes are not likely to be drawn from the blue curve in Figure 1A—requires either deeper sampling (i.e. more replicates and/or more guides targeting each gene) or a more severe phenotype. In fact, this is reflected in the MAGeCK results for the four TKO cell lines tested: the GBM and RPE1 cell lines were screened with a 90k library and MAGeCK yielded 586 and 489 hits, respectively, while the HeLa and HCT116 lines were screened with a 177k library and MAGeCK yielded 718 and 905 hits – on average, ~50% more hits using the larger library. The sequence-optimized 180k gRNA library used by Wang *et al*. has a lower proportion of non-performing guides, resulting in substantially improved sensitivity for both BAGEL and MAGeCK, though BAGEL still identifies ~50% more hits in each screen.

## Conclusions

The ability to perform accurate, saturating genetic perturbation screens in human cell lines could transform molecular genetics in the coming years. To maximize potential—and to avoid pitfalls similar to the costly false starts encountered in the RNAi field—rigorous analytical methods must be applied that are able to effectively discriminate true hits from false positives. While data suggests that off-target effects in CRISPR-Cas9 pooled library screens are much less of a concern than with RNAi, the variable effectiveness of early reagent pools makes it important that analytical methods are able to detect subtle phenotypes. BAGEL accurately models the wide variability in phenotype shown by reagents targeting known essential genes, enabling the sensitive and precise identification of fitness genes, even under conditions of suboptimal data quality.

## Availability and requirements

Project name: bagel-for-knockout-screens

Project home page: http://bagel-for-knockout-screens.sourceforge.net/

Operating system(s): platform independent

Programming language: Python

Licensing: [to be determined]

No restriction for non-academic use.

TKO screen data are available at http://tko.ccbr.utoronto.ca/

## Authors’ contributions

TH developed the algorithm and wrote the software; TH and JS wrote and edited the manuscript. All authors read and approved the final manuscript.

## Acknowledgements

We would like to thank Megha Chandrashekhar, Michael Aregger, Zachary Steinhart, and Stephane Angers for acquiring the data for this study.

